# Probe Efficient Feature Representation of Gapped K-mer Frequency Vectors from Sequences using Deep Neural Networks

**DOI:** 10.1101/170761

**Authors:** Zhen Cao, Shihua Zhang

**Affiliations:** NCMIS, CEMS, RCSDS, Academy of Mathematics and Systems Science, CAS, Beijing 100190; School of Mathematical Sciences, University of Chinese Academy of Sciences, Beijing 100049, China..

**Keywords:** Bioinformatics, machine learning, gapped k-mer, deep neural network, transcription factor binding site pediction

## Abstract

How to extract informative features from genome sequence is a challenging issue. Gapped k-mers frequency vectors (gkm-fv) has been presented as a new type of features in the last few years. Coupled with support vector machine (gkm-SVM), gkm-fvs have been used to achieve effective sequence-based predictions. However, the huge computation of a large kernel matrix prevents it from using large amount of data. And it is unclear how to combine gkm-fvs with other data sources in the context of string kernel. On the other hand, the high dimensionality, colinearity and sparsity of gkm-fvs hinder the use of many traditional machine learning methods without a kernel trick. Therefore, we proposed a flexible and scalable framework gkm-DNN to achieve feature representation from high-dimensional gkm-fvs using deep neural networks (DNN). We first proposed a more concise version of gkm-fvs which significantly reduce the dimension of gkm-fvs. Then we implemented an efficient method to calculate the gkm-fv of a given sequence at the first time. Finally, we adopted a DNN model with gkm-fvs as inputs to achieve efficient feature representation and a prediction task. Here, we took the transcription factor binding site prediction as an illustrative application. We applied gkm-DNN onto 467 small and 69 big human ENCODE ChIP-seq datasets to demonstrate its performance and compared it with the state-of-the-art method gkm-SVM. We demonstrated that gkm-DNN can not only improve the limitations of high dimensionality, colinearity and sparsity of gkm-fvs, but also make comparable overall performance compared with gkm-SVM using the same gkm-fvs. In addition, we used gkm-DNN to explore the representation power of gkm-fvs and provided more explanation on how gkm-fvs work.

## 1 Introduction

IT is still a major challenge to study the function of primary DNA sequences in non-coding regions [1], [2]. The regulatory elements located in these regions usually contain several transcription factor binding sites (TFBSs), whose activities regulate multiple biological processes such as gene expression [3]. More importantly, some genetic variations in regulatory elements cause hereditary disease, which attracts more attention [4]. In the past decade, several large-scale projects such as Encyclopedia of DNA Elements (ENCODE) and Roadmap Epigenomics Program have been launched to generate and profile large amounts of genome-wide data to figure out the mechanisms behind these regulatory processes [5], [6].

In this background, using statistical and machine learning methods to study the regulatory elements is a promising paradigm. For example, Wang et al. [7] used an expectation maximization (EM) based method to analyze position weight matrix (PWM) for 457 ENCODE ChIP-seq data sets of 119 different transcription factors. They profiled the sequence features and chromatin structure around the corresponding TFBSs. Liu et al. [8] integraed oligomers of short length (known as l-mers) and six DNA local parameters to identify enhancers along with their intensity.

Ghandi et al. [9] used gapped k-mer (oligomers of length l with l – k non-informative positions) frequency vectors and support vector machines (gkm-SVM) to precisely predict the TFBSs. They also successfully used the scores of gkm-SVM to predict the impact of regulatory variants from DNA sequence [10]. More recently, Kelley et al. [11] used one-hot coding format of DNA sequences and then applied deep convolutional neural networks (CNNs) to predict the chromatin accessibility of 164 cell types, and interpret some disease-related single nucleotide polymorphisms.

Yet, how to extract informative features from genome sequence is still a primary challenging issue despite the great success in studying the regulatory elements. The core of all above approaches is extracting features from genome sequences, while all of them have pros and cons. The position weight matrix (PWM) approach is close to the biological nature (e.g., motifs), but requires large amounts of data to determine appropriate scoring thresholds [12]. The onehot coding format of DNA sequences resolves limited information explicitly, which puts the heavy pressure on model training [11], [13], [14]. As a supplement, using the frequencies of all l-mers to form feature vectors (l-mer frequency vector, abbreviated as lmer-fv) can resolve most of the information before training and prediction, while they becomes sparse and unstable as l increases [15]. Compared to lmer-fvs, gapped k-mer frequency vectors (abbreviated as gkm-fvs) are not only close to the biological nature, but also more robust [15]. Therefore, gkm-fv based methods have been used as benchmark methods for many bioin-formatics problems [11], [14], [16].

However, there still exist limitations to be addressed when using gkm-fvs as features. The dimensions of gkm-fvs grow exponentially as either l or k increases, which quickly become intractable. Besides, gkm-fvs are somehow sparse and highly collinear due to the inner structure of gkm-fvs [15]. The popular method gkm-SVM adopts support vector machines with kernel tricks to tackle the problems [9], [17]. However, gkm-SVM cannot quickly deal with large amount of data due to inefficient computation of a large-scale kernel matrix. And it is unclear how to combine gkm-fvs with other data sources in the context of a string kernel. The shortcomings of kernel tricks may hinder the advantage of gkm-fvs. Another consideration is to directly use gkm-fvs to solve several issues. First, it is not a trivial task to calculate the gkm-fvs for large amount of sequences especially in the absence of a quick method to handle this problem. Second, the high dimensionality and colinearity imply the structural redundance of raw gkm-fvs, which needs a more efficient representation. Third, the colinearity and sparsity of gkm-fvs hinder the use of many traditional machine learning methods such as logistic regression. Thus, efficient methods are still needed to make full use of gkm-fvs.

In the past few years, the fast development of deep neural networks (DNNs) attracts great attentions in bioinformatics community due to several reasons. First, the rapid accumulation of diverse biological data fits in with the usability of DNN models. Second, the use of graphics processing unit (GPU) makes the training process of DNNs extremely faster than before [18], [19]. Third, it is easier to train models with the help of DNN specific technologies such as dropout and batch normalization [20], [21]. Therefore, DNNs have achieved state-of-the-art performance in a wide range of applications such as image classification and speech recognition [22], [23]. In addition, bioinformatics community has also successfully applied DNNs to the predictions of transcription factor binding sites, genome variants, gene expression and so on [13], [14], [24]. The power of DNN in handling extremely large-scale datasets and learning hierarchical non-linear data representation enables it has the potential to make full use of gkm-fvs.

In this study, we proposed a flexible and scalable method gkm-DNN to achieve feature representation and prediction from high-dimensional gkm sequence features using DNN (Fig. 1). We first proposed a more concise version of gkm-fvs, which significantly reduce the dimensions of raw gkm-fvs. Then we implemented an efficient method to calculate the gkm-fv of a given sequence at the first time (Fig. 1A). Next we took the gkm-fvs as input for DNN to achieve a prediction task (Fig. 1B). We took the TFBS prediction as an illustrative example, which is a widely studied problem. We demonstrated that gkm-DNN can not only improve the potential limitations of high dimensionality, colinearity and sparsity of gkm-fvs, but also make comparable overall performance compared with gkm-SVM using the same gkm-fvs.

**Fig. 1.**
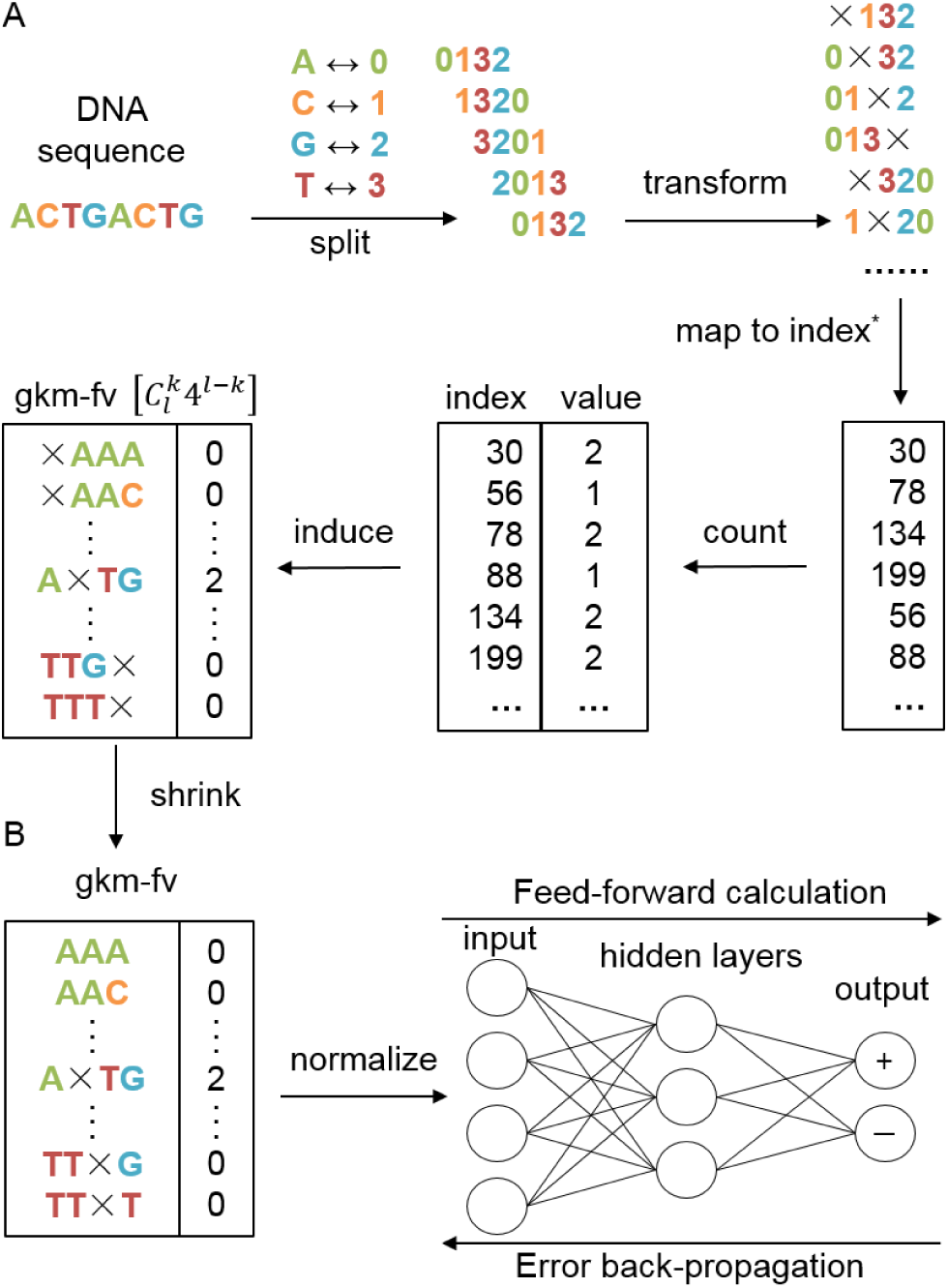
The key components of constructing a gkm-DNN prediction model. (A) An efficient method to calculate the gapped k-mer frequency vector (gkm-fv). (B) Illustration of a DNN model. After normalization, the gkm-fvs are taken as input for a multi-layer perceptron model. This model is trained using the standard error back-propagation algorithm and mini-batch stochastic gradient descent method. ^*^Note we have adopted an efficient ‘map to index' calculation trick to do it (see Materials and Methods section).

## 2 Materials and Methods

### 2.1 Datasets

We downloaded all the 467 ENCODE ChIP-seq datasets used in [9] from http://www.beerlab.org/gkm-SVM/ to demonstrate the performance of gkm-DNN and compared it with gkm-SVM. All positive and negative sequences are the same as used in [9]. Each dataset contains at most 5000 positive sequences and the same number of negative sequences.

We also collected 69 high-quality datasets of larger size for further tests from ENCODE [5]. To ensure the quality of data, we downloaded all the datasets in the format of optimal idr thresholded peaks from the K562 cell lines. We only retained peaks with q-value < 0.01 and selected datasets with more than 25000 peaks. Given a dataset, we sort peaks first by ‐log(q-value), and then by its count in descending order. We only kept the top 20000 peaks for further analysis since there are always a lot of noises in ChIP-seq datasets [25]. If a peak is shorter than 301bp, we extended this peak from its midpoint to 301bp. If a peak is longer than 601bp, we cut down the peak to 601bp centered at the raw midpoint. After processing, we finally got 69 high-quality datasets. Each dataset contains 20000 positive samples.

Given positive samples, we also generated the same number of negative samples. We randomly sampled sequences of approximately the same length of positive samples from the whole genome. The randomly selected sequences were dropped if they have overlap with optimal idr thresholded peaks.

### 2.2 Calculating the Concise gkm-fvs

Oligomers of fixed length l are commonly known as l-mers. Gapped k-mers are l-mers with l – k noninformative positions (Supplementary Fig. S1). In other words, for gapped k-mers, there are two parameters: word length l and matched position k (k < l, hence there are l – k gaps). Given l and k, the frequencies of all gapped k-mers form a vector of length, which is called gapped k-mer frequency vector (gkm-fv) in this paper. Different from the kernel computation of gkm-SVM, our method directly uses the gkm-fvs as input. However, it is not a trivial task to calculate the gkm-fvs of hundreds of thousands of DNA sequences. Almost all methods use kernel tricks to skip its direct calculation. To the best of our knowledge, there is no quick program to directly calculate the gkm-fvs. For the first time, we implemented an efficient method to calculate it.

Given a single strand DNA sequence s, we split it into length(s)-l+1 l-mers and then transformed the l-mers to gapper k-mers. We turned the gapped k-mers into a quaternary representation (A→0, C→1, G→2, T→3), which induces their positions in the ordered gkm-fv to be a decimal integer. For example, G×CA→2× 10→(gap position 1, 210 of base 4)→(gap position 1, 36 of base 10) →1*43+36 →100 (G×CA→100 for short). The computing using the quaternary representation is efficient in both time and space, which skips the calculation of string matching. Then we counted the frequencies of gapped k-mers and automatically got the desired gkm-fvs. Besides, we also implemented a slightly faster method for counting relatively short gkm-fvs. We first counted the lmer-fvs using the above quaternary representation and then transformed the lmer-fvs to gkm-fvs using a matrix multiplication strategy [15].

To handle the high dimensionality of raw gkm-fvs, we proposed a concise version of gkm-fvs which deals with gaps at the ends. Since the gaps at the ends provide little information, we degenerated the gapper k-mers with gaps at the ends to a shorter gapped k-mer or a simple k-mer (Supplementary Fig.1). For example, ×A×G and A×G× both shrink to A×G. This step significantly reduces the dimensions of raw gapped k-mers (Fig. 2A and supplementary Fig.3A) and simplifies all the calculations of gkm-DNN.

**Fig. 2.**
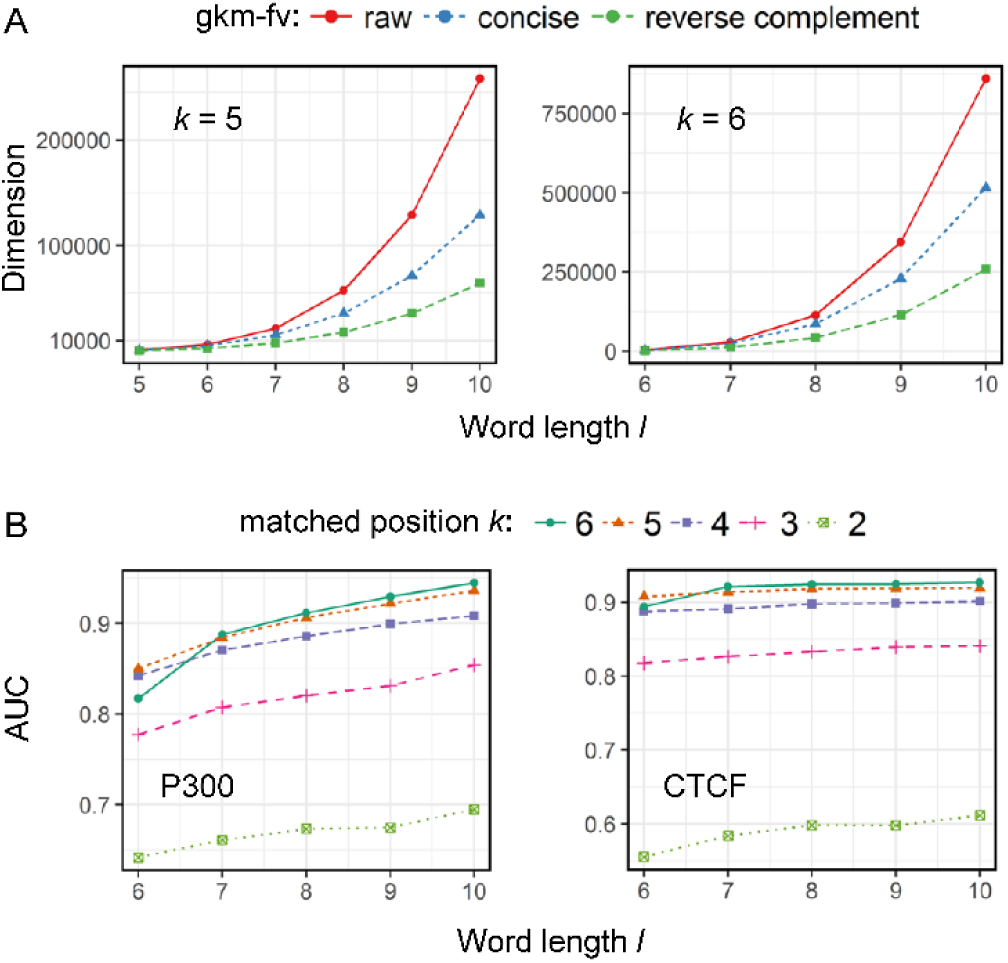
Performance comparison between different gkm-fvs. (A) Dimensions of different gkm-fvs in three situations: raw (red), concise version (blue), and concise version using reverse complement DNA strand (green). Matched position k = 5 for left panel and k = 6 for right panel. (B) AUCs of two representative datasets (P300 and CTCF in GM12878 cell line) with respect to different gkm-fvs (concise version using reverse complement DNA strand) are shown. Given word length l and matched position k, the best model is trained automatically. Each point represents the AUC of a type of gkm-fvs.

In this paper, we considered the double strands DNA sequences. So we counted gapped k-mers in DNA double strands. Instead of counting exact gapped k-mers, we also counted their reverse complements and remove redundant gapped k-mers. For example, two gapped k-mers A××CTG and CAG××T are reverse complements to each other and have exactly the same frequencies. Thus, we only kept one of them. This step not only reduces about 50% of the input sizes (Fig. 2A and Supplementary Fig.3A), but also well captures the double strands information.

We normalized the frequency vectors before training the DNNs. Given a DNA sequence, we divided its gkm-fv by the length of the sequence. Then we multiplied the frequencies with min(4^*k*^ /2,128) to reduce the floating point error caused by GPU calculation.

### 2.3 gkm-DNN Model

Here we adopted a multi-layer feedforward neural network model to achieve the prediction task with the gkm-fv of a sequence as an input feature vector (Fig. 1B). It contains one input layer, one or multiple hidden layers and one output layer. All the layers are fully-connected. The feedforward calculation is a linear transformation with an activation function (Fig. 1B). Specifically, given the values of layer m-1, the values of layer m is calculated as:

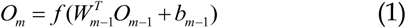

where *O*_*m*–1_ is the values of layer m-1, {*W*_*m*–l_,*b*_*m*–1_} are the weights and the bias associated with layer m-1 that need to be learned. For the hidden layers, *f*(·) is the RELU activation function. The sigmoid activation function (or softmax function) is applied to the output unit for the classification purpose:

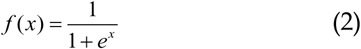

The cross entropy is used as the loss function for training, namely,

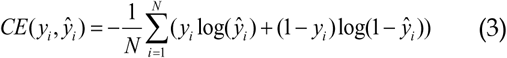

where *ŷ*_*i*_ is the output of an input sequence, *y*_*i*_ = 1 for a positive sequence, *y*_*i*_ = 0 for a negative sequence and *N* is the number of all training samples. Given the loss function and hyper-parameters (see below), the model is trained using the standard error back propagation algorithm and mini-batch stochastic gradient descent method. We also used two tricks namely dropout and early stopping (Supplementary Materials and Fig. S2). Passing all the training data through the model once is an epoch. Training is stopped when it hits the maximal training epoch given in advance.

### 2.4 Setup of gkm-DNN

Given the training sequences, we first calculated the gkm-fvs and then normalized them. Then we randomly split the total training set into a smaller training set and a validation set ( 1 / 8 of the total training set in this paper). Given different groups of hyper-parameters (Supplementary Table S1 and S2), the best model was chosen according to the cross entropy on validation set.

Given a test DNA sequence, we calculated the gkm-fv using corresponding l and k and normalized it. Next we used the best model to do prediction. The output is an estimated posterior probability on how likely this sequence is a positive one (i.e., how likely the transcription factor binds to this sequence).

There are some hyper-parameters for gkm-DNN. For all models, we always applied dropout to the last layer. The dropout rate was set as {10%, 20%, 30%} for further discussion [20]. The depth and width are different for datasets of different sizes. We use P300 and CTCF datasets to test the performance of different gkm-fvs (Supplementary Table S1) and set the number of hidden nodes as {100, 200, 400, 700} here. For the 467 small ENCODE ChIP-seq datasets (max sample size: 10000), we only used one hidden layer and set the number of hidden nodes as {100, 400, 700} for further analysis. For these 467 ChIP-seq datasets, we trained two groups of models for further discussion. First, we trained models using the default gkm-fvs (l = 7, k = 5, 7680 dimensions). Second, we trained models using longer gkm-fvs (l = 8, k = 6, 43168 dimensions) to the best of our computing power. In order to eliminate the effect of small sample sizes, we retrained models with the best hyper-parameters using all training data (validation set included) on the basis of the second part. For the 69 high-quality datasets (sample size: 40000), we set the number of hidden nodes as 700 and the number of hidden layers as {1, 2, 3} for further analysis. We trained models using the default gkm-fvs (l = 7, k = 5). More details and motivations about hyper-parameters of gkm-DNN can be seen in the Supplementary Materials, Tables S1, S2 and S3.

In this paper, we applied several practical ways to accelerate the training (Supplementary Materials). For input, we saved our data in a binary format in advance, which greatly reduces the time of input and output. For calculation, we used GPU (one NVIDIA GTX 1080) to train the model with half-precision floating-point.

### 2.5 Setup of gkm-SVM

In this paper, we used both the gkm-SVM R package and LS-GKM to implement gkm-SVM [17], [26]. On 467 small datasets, we trained gkm-SVM with three different gkm-fvs for comparison (l = 7, k = 5 and l = 8, k = 6 and l = 10, k = 6). On 69 big ChIP-seq datasets, gkm-SVM R package is not able to handle all the training sets. So we randomly selected 10000 sequences from the whole training sets and ran gkm-SVM R package with l = 10, k = 6. For comparison, we also ran LS-GKM with l = 7, k = 5 and l = 10, k = 6 using the whole datasets. Other parameters are set as defaults.

We applied several practical ways to accelerate the training of gkm-SVM. For gkm-SVM R package, we first calculated the full kernel matrix and then used the crossvalidation mode. This step greatly reduces the repetitive computation of a kernel matrix. In addition, the gkmSVM tasks were also parallelized using R language. For LS-GKM, we used all the four cores (four threads) and 10G cache memory, which makes full use of our machine.

### 2.6 Performance Assessment

We adopted multiple standard measures to comprehensively evaluate gkm-DNN and gkm-SVM. The receiver operating curves (ROC) and precision-recall curves (PRC) are two typical graphical plots that illustrate the classification ability of a binary classifier system as its discrimination threshold is varied [27], [28]. We used the area under the ROC and PRC curves (AUC and AUPR). However, many studies have illustrated the drawbacks of ROC and AUC [29], [30]. The areas under the curves essentially only assess the ability of ranking samples. One rarely uses AUC in a practical classifier because only a couple of points on the curves are useful [30]. For a typical prediction task such TFBS prediction, it is very important to give a solid binary prediction [31]. Thus, we also adopted four widely used performance measures including

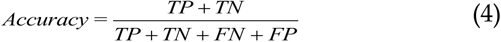

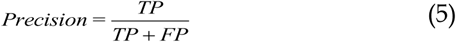

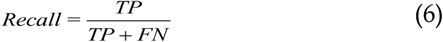

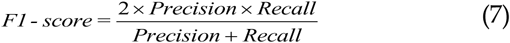

where TP, FP, TN and FN represent true positives, false positives, true negatives and false negatives, respectively. In addition, we used the standard 5-fold cross-validation procedure to get the mean of above measurements.

### 2.7 Training DNN with Different Size of Samples

We also trained gkm-DNN models with different size of training samples using the 69 high-quality datasets. For each dataset and each cross-validation fold, given the same 4000 validating samples, we used training samples of equal size (1×), twice (2×), four times (4×) and seven times (7×) to train the gkm-DNN models and calculated the AUCs and accuracies on the test sets. To strictly control the quality of data, 1× samples are contained in 2× samples, 2× samples are contained in the 4× samples and so on. Note that using 7× samples is the same as using all training samples as above. The maximum training epochs were set as 210, 105, 53 and 30 for 1×, 2×, 4× and 7× training samples. All other hyper-parameters were set as the same one.

### 2.8 Implementation of gkm-DNN

We used Scala and R to calculate the gkm-fvs of DNA sequences. The implementation using Scala with GPU is approximately seven times faster than using R. We adopted DL4J framework (deep learning for Java, version 0.8.0) to train neural networks. All the source codes are available at http://page.amss.ac.cn/shihua.zhang/.

## 3 Results

### 3.1 Evaluation of Different gkm-fvs

To address the high dimensionality of raw gkm-fvs, we first proposed a more concise version of gkm-fvs, which have multiple advantages. First, this step reduces the dimensions of raw gkm-fvs with little information loss. The rates of reductions vary from 15% to 80%, which greatly reduces the computational burden (Fig. 2 and Supplementary Fig. S3). Second, the concept of gkm-fvs becomes more coherent because when k is the same, a gkm-fv with shorter l is always a subset of a gkm-fv with longer l. Hence a gkm-fv can be seen as an extension of the corresponding kmer-fv. In addition to the concise gkm-fvs, we also used the double strands DNA information to reduce approximately half of the input dimensions (Fig. 2 and Supplementary Fig. S3A). All these efforts simplified the gkm-fvs from structure and made us able to use some longer gkm-fvs with less computation.

For running gkm-DNN, the first step is to select proper l and k to determine gkm-fvs. Here, we used two representative datasets P300 and CTCF from GM12878 cell line to evaluate the effects of gkm-fvs in terms of l and k. P300 is a coactivator whose binding events usually mark the enhancer. Its underlying binding sites may be comprehensive but complex. On the other hand, CTCF is reported to have specific and long binding sites. Therefore, the two datasets could guide us to select proper gkm-fvs.

Generally, different gkm-fvs in terms of l and k significantly influence the performance of gkm-DNN. Given l, the larger the k is, the better the performance is (Fig. 2B). When k >= 4, the performances are significantly better than that of k = 2 or 3 (Fig. 2B). Thus, very small k cannot resolve enough information from DNA sequences. On the other hand, given k, the longer the l is, the higher the AUC is (Fig. 2B). However, the results of lmer-fv (l = 6) is worse than that of gkm-fvs (l = 6, k = 5), implying the necessity of proper gaps (Fig. 2B). The results in terms of AUPR, accuracy and F1-score show similar performance (Supplementary Fig. S3B). Thus, we noted that large l and proper k may resolve more information from raw DNA sequences directly.

On the other hand, the performances of P300 and CTCF datasets are different. The gkm-fvs with longer l greatly improves the prediction of CTCF binding sites while the improvement for P300 is quite small. As for P300, there seems to be great space for improvement while the longer gapped k-mers alone are not enough. For these problems, integrating more and other types of data or using other machine learning frameworks may help. Thus, we finally chose l = 7, k = 5 (7680 dimensions) as default parameters for gkm-DNN due to the computational efficiency and tried to integrate more data in the following experiments.

### 3.2 Performance on Human ENCODE ChIP-seq Datasets

Here we applied gkm-DNN onto the 467 human ENCODE ChIP-seq datasets and compared it with gkm-SVM 2.0. Combination of gkm-fvs and DNN demonstrate strong potentials. At first, we used the default gkm-fvs of gkm-DNN (l = 7, k = 5) for both methods (Fig. 3A). The two methods show very competitive performances in terms of AUC and F1-score (Fig. 3A). To further confirm the potential of combination of gkm-fv and DNN, we also trained models using longer gkm-fvs (l = 8, k = 6, 43168 dimensions), which reaches the ultimate capacity of our equipment computing power. The two methods both improve the results and also show competitive performances in terms of AUC and F1-score when using longer gkmfvs (Fig. 3A).

**Fig. 3.**
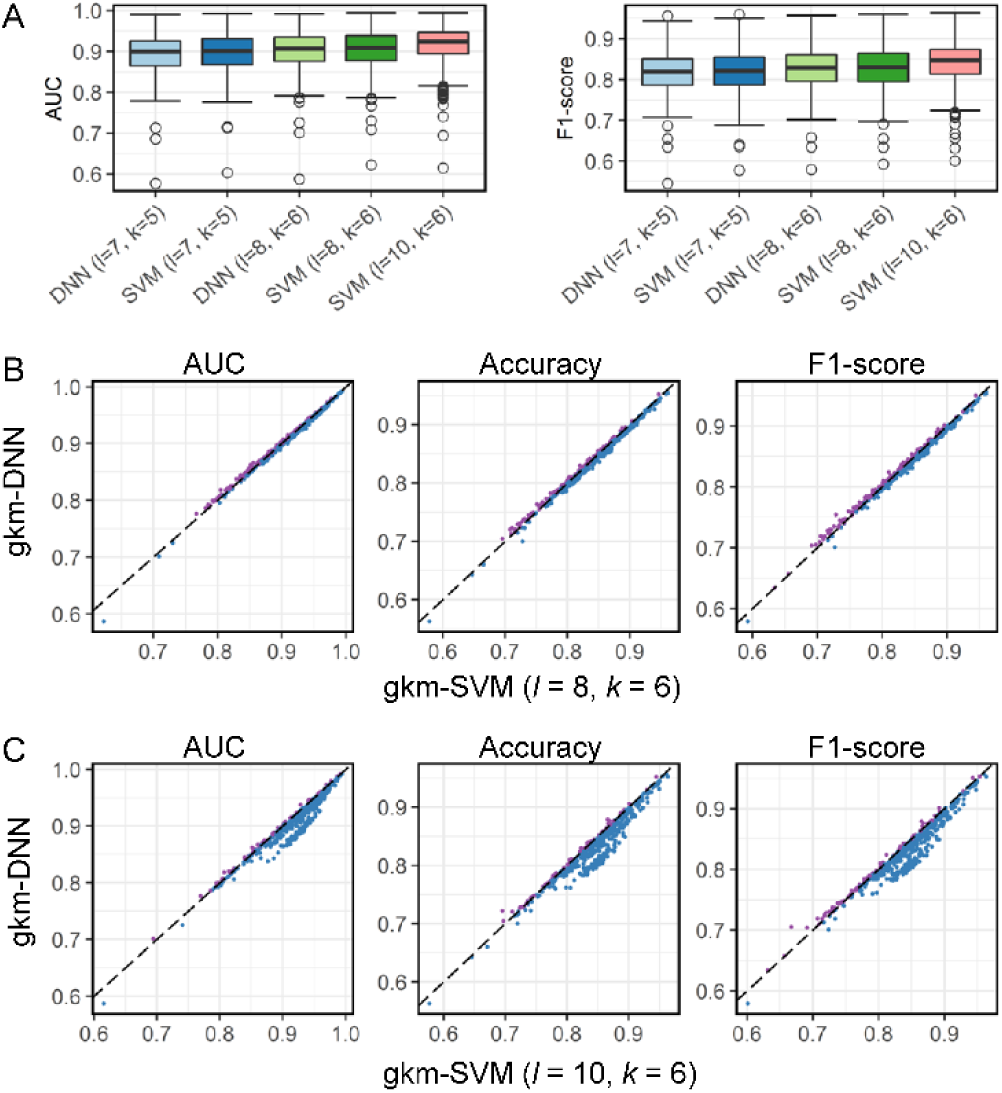
Performance comparison between gkm-DNN and gkm-SVM on 467 small ChIP-seq datasets. (A) Box plots of AUC and F1-score for different models. The bottom, top, and middle bands of the boxes indicate the 25th, 75th, and 50th percentiles, respectively. Whiskers extend to the most extreme data points no more than 1.5 interquartile range from the box. (B) and (C): scatter plots in terms of AUC, accuracy and F1-score. Each point represents the prediction results of gkm-DNN (y-coordinate) and gkm-SVM (×-coordinate) of a dataset. Points above and below the line y = × are in purple and blue respectively. (B) Comparison between gkm-DNN (l = 8, k = 6) and gkm-SVM (l = 8, k = 6). (C) Comparison between gkm-DNN (l = 8, k = 6) and gkm-SVM (l = 10, k = 6).

To further compare the two methods, we analyzed the results on each of the 467 datasets. The two methods show almost the same performances in terms of AUC, accuracy and F1-score on all 467 datasets (Fig. 3B). But, gkm-DNN usually has higher recalls and lower precisions (Supplementary Fig. S4A). This may be caused by the difference between structural risk minimization (SVM) and empirical risk minimization (DNN). Surprisingly, the number of parameters (>4, 316, 800) is five hundred times bigger than the size of training sample (8000), while gkm-DNN models are not over-fitted due to multiple normalization strategies such as early stopping.

We also compared gkm-DNN (l = 8, k = 6) with gkm-SVM using its default parameters (l = 10, k = 6). Currently, it is hard for gkm-DNN to use large amount of features (l = 10, k = 6, 258496 dimensions) on hundreds of datasets due to our poor equipment. More importantly, it is not a good idea to include too many features as the sample size is relatively small. gkm-SVM indeed outperforms gkm-DNN in terms of AUC, AUPR, accuracy and F1-score in most cases (Fig. 3A, Fig. 3C and Supplementary Fig. S4B).

### 3.3 Performance on High-quality Datasets of Larger Size

Here we applied gkm-DNN onto 69 high-quality datasets with relative larger sizes (40000 samples per dataset) and compared it with gkm-SVM and LS-GKM. Using more data, gkm-DNN achieved comparable AUCs with gkm-SVM (mean differences of AUCs 0.00002, Figs. 4A and 4B). gkm-DNN got higher AUCs on 65% of the datasets (45 out of 69), indicting the importance of more training data. On the other hand, gkm-DNN got distinct better accuracies and F1-scores than gkm-SVM (Figs. 4A and 4B). We also observed the phenomenon when comparing gkm-SVM and LS-GKM (Fig. 4A). So the small training samples may result in the instability of the desired classifier, which further strengthen the importance of using more data.

**Fig. 4.**
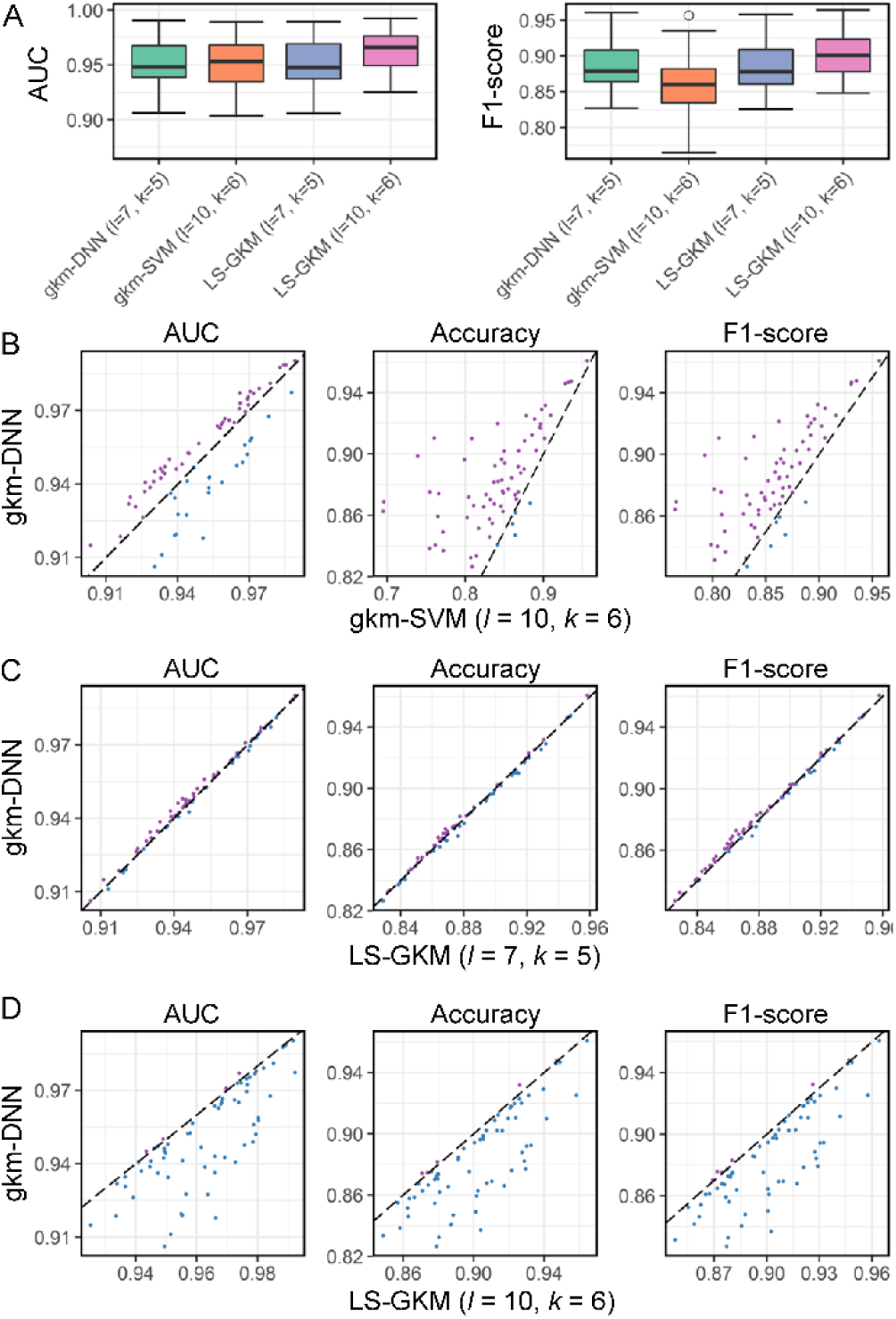
Performance comparison between gkm-DNN and gkm-SVM on 69 high-quality ChIP-seq datasets. (A) Box plots of AUC and F1-score for different models. The bottom, top, and middle bands of the boxes indicate the 25th, 75th, and 50th percentiles, respectively. Whiskers extend to the most extreme data points no more than 1.5 interquartile range from the box. (B), (C) and (D): scatter plots in terms of AUC, accuracy and F1 score. Each point represents the prediction results of gkm-DNN (y-coordinate) and gkm-SVM/LS-GKM (×-coordinate) of a dataset. Points above and below the line y = × are in purple and blue respectively. (B) Comparison between gkm-DNN (l = 7, k = 5) and gkm-SVM using less training data (l = 10 k = 6). (C) Comparison between gkm-DNN (l = 7, k = 5) and LS-GKM (l = 7, k = 5). (D) Comparison between gkm-DNN (l = 7, k = 5) and LS-GKM (l = 10, k = 6).

For gkm-DNN and LS-GKM, the results on these big datasets show very similar performance to those on the small datasets. When using the same features (l = 7, k = 5), the two methods are comparable in terms of AUC, accuracy and F1-score (Fig. 4C); while gkm-DNN usually has higher recalls and lower precisions (Supplementary Fig. S5B). When LS-GKM uses longer gapped k-mers (l = 10, k = 6), it can achieve higher AUC, AUPR, accuracy and F1-score (Fig. 4D and Supplementary Fig. S5C).

In conclusion, more data and longer gapped k-mers are both important. As suggested by our experiments, all three methods have their own pros and cons. gkm-DNN can quickly deal with large amount of data at the cost of using relatively shorter gapped k-mers. gkm-SVM 2.0 is fast while it can only deal with small or medium datasets. LS-GKM is able to deal with both large datasets and longer gapped k-mers. However, LS-GKM is quite slower than the above two methods (see below). Generally, choosing a proper method depends on the specific circumstance.

### 3.4 Computational Efficiency

Besides feature representation and prediction accuracy, we aimed to develop a fast method which can handle large-scale datasets. To verify the efficiency of gkm-DNN, we tested the average runtime of gkm-DNN and gkm-SVM under default parameters using our desktop computer (Supplementary Table S4). On small datasets (sample size 10000), gkm-DNN and gkm-SVM 2.0 are both fast enough while gkm-DNN is a little faster. However, LS-GKM is much slower than the two methods (Fig. 5A). The reason is that LS-GKM does not presave the kernel matrix and does many repetitive computation. On big datasets (sample size 40000), the gkm-SVM 2.0 package cannot handle this case because it needs too much memory. Moreover, gkm-DNN is about 37 times faster than LS-GKM (Fig. 5A).

**Fig. 5.**
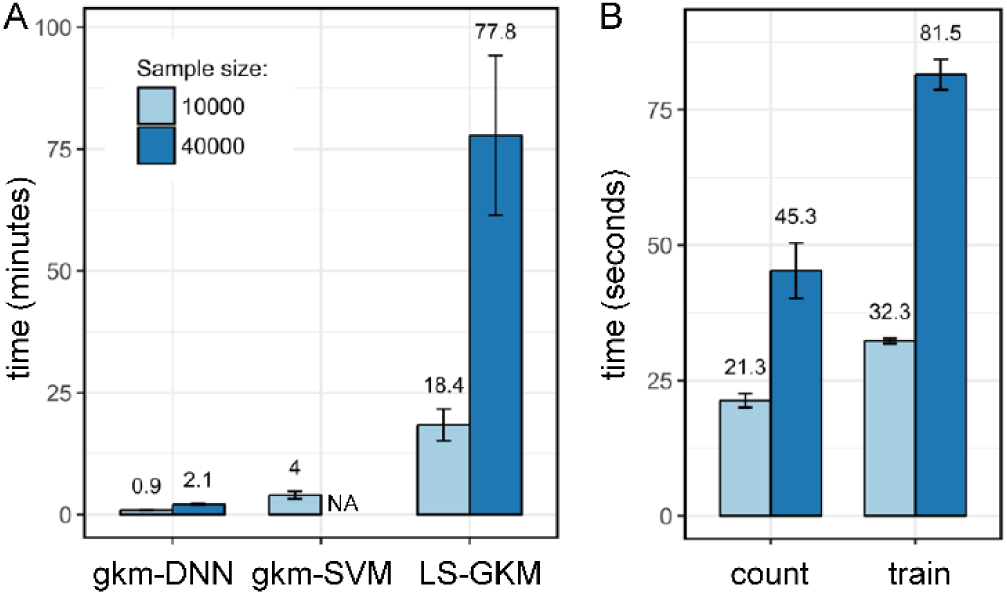
Computational efficiency of gkm-DNN. (A) Comparison of computing time between gkm-DNN, gkm-SVM 2.0 and LS-GKM. The average runtime of both methods for one dataset using a desktop computer is presented. (B) Detailed average runtime of gkm-DNN on small and big datasets. Training gkm-DNN consists of calculating gkm-fvs (count) and training DNN (train).

More importantly, the runtime of gkm-DNN is proportional to the sample size. We explored the runtime of its two components, namely calculating gkm-fvs (along with saving gkm-fvs into binary format) and training DNN (Fig. 5B). The parts can be naturally paralleled at the sample level. Thus, the runtime of each part should be proportional to sample size. We confirmed this by comparing the average runtime of gkm-DNN on small and big datasets (Fig. 5B). On big datasets, the running time is faster than expected because the program makes more use of our desktop computer.

### 3.5 gkm-DNN has Great Superiority with More Data

We observed that the performance on big datasets are greatly better than that on small datasets (e.g., the average AUCs are 0.952 and 0.903 for gkm-DNN respectively) (Figs. 3 and 4). Therefore, we believe that larger sample size is also important in real applications. In this situation, gkm-DNN have great superiority because it can quickly make full use of large amount of data as shown above.

We trained gkm-DNN based on the 69 high-quality datasets using different number of training samples (Materials and Methods). We first observed that models trained from more data have overall higher AUCs and F1-scores than models trained from less data (Fig. 6). Furthermore, for each dataset, the results of gkm-DNN are always better when using more data (Supplementary Fig. S6). In short, gkm-DNN can achieve an improvement using more training samples. With rapid development of biological technology, more DNA sequence data will emerge and gkm-DNN will definitely benefit from these large-scale data.

**Fig. 6.**
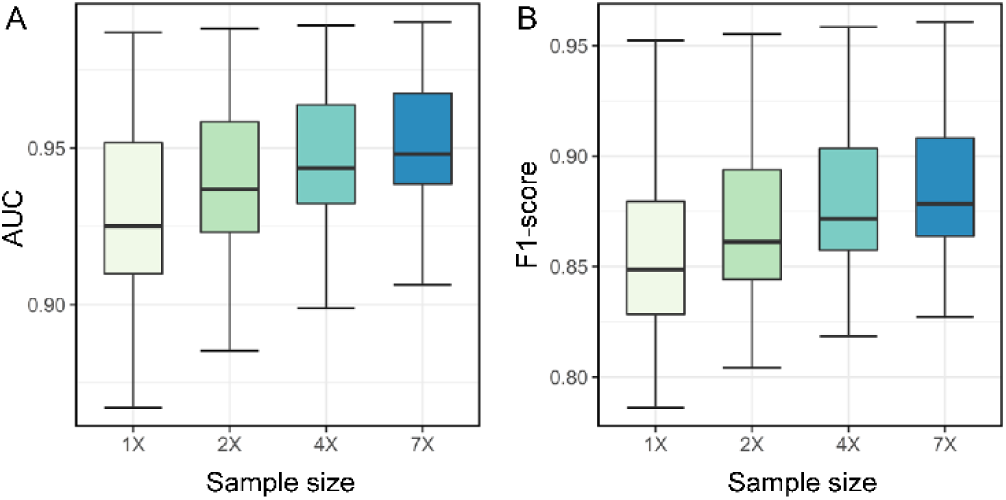
Performance comparison of gkm-DNN trained using different sample sizes in terms of AUC and F1-score. For each dataset and each cross-validation fold, given the same validation set, training sets of equal size (1×), twice (2×), four times (4×) and seven times (7×) were used to train gkm-DNN. For each dataset, average AUC (A) and F1-score (B) were calculated on the test sets for five-fold cross-validation.

### 3.6 Interpretation of gkm-DNN

Here, we attempted to elucidate the learned neural networks and figure out how gkm-DNN works using the 69 high quality datasets. For inputs, we can view the gkm-fvs as a type of auto-encoder of lmer-fvs (Supplementary Fig. S7). In practice, gkm-SVM uses gkm-fvs or estimated lmer-fvs based on gkm-fvs as input (Supplementary Fig. S7). We can view the input of gkm-DNN as a denoising version of lmer-fvs, which makes the classifier easy to learn.

Moreover, using hidden layers can computationally improve the potential limitations of high dimensionality, colinearity and sparsity of gkm-fvs. Compared to 7680 raw features, the hidden layers have at most 700 nodes (Supplementary Fig. S8A), which forces the models to learn the most representative features. Furthermore, the learned activation values of hidden nodes are not redundant. Given a trained model and all training samples, we calculated the condition numbers of the activation values of the last hidden layer. The condition numbers are all very small (Supplementary Fig. S8B), indicating that the features extracted by hidden layers are not collinear at all. In addition, the activation values of the last hidden layer are denser than the gkm-fvs. Although these values contain zeros due to the RELU activation function, the ratio of zeros is distinctly lower than that of gkm-fvs (Supplementary Fig. S8C).

The hidden layers learn useful patterns for the classification task. Here we took one learned model as an example. For the last hidden layer, we clustered both the 700 hidden nodes and the 32000 training samples using the activation values (Supplementary Fig. S8D). There are clear clusters in the heatmap, which distinguish a fraction of training samples from others (Supplementary Fig. S8D). We also employed t-distributed stochastic neighbor embedding (t-SNE) to transform the 700 features into two representative directions [32]. The two directions can separate many positive samples from negative samples distinctly (Supplementary Fig. S8E). Moreover, the distribution of negative samples are more disperse than that of positive ones. The negative samples are randomly selected from the whole genome, implying that they may contain no much strong signals. In contrast, the positive samples may contain some underlying patterns (e.g., motifs). Thus, gkm-DNN distinguishes positive samples using the combination of some weak patterns from the negative ones.

### 3.7 gkm-DNN Provides More Insights into gkm-fvs

We further found that the models with more hidden layers are marginally worse than those with less hidden layers (Fig. 7A), indicating that deeper models don't enhance the performance of gkm-fvs. For further analysis, we took the CTCF dataset (from K562 cell line with 40000 samples) as an example to test more complex models. First, adding more hidden layers slightly decreased the accuracies (Fig.7B). To rule out the possibility of gradient vanishing and optimization difficulty of deep models, we added batch normalization (BN) or residual connection in the previous models [21], [33]. However, all these complex models perform significantly worse than the simplest one (Fig. 7B), which is quite counterintuitive. The above observation points that the neural network models are easily over-fitted using the gkm-fvs as inputs. Moreover, we suspected that the gkm-fvs contains few structured information to be mined by neural networks.

**Fig. 7.**
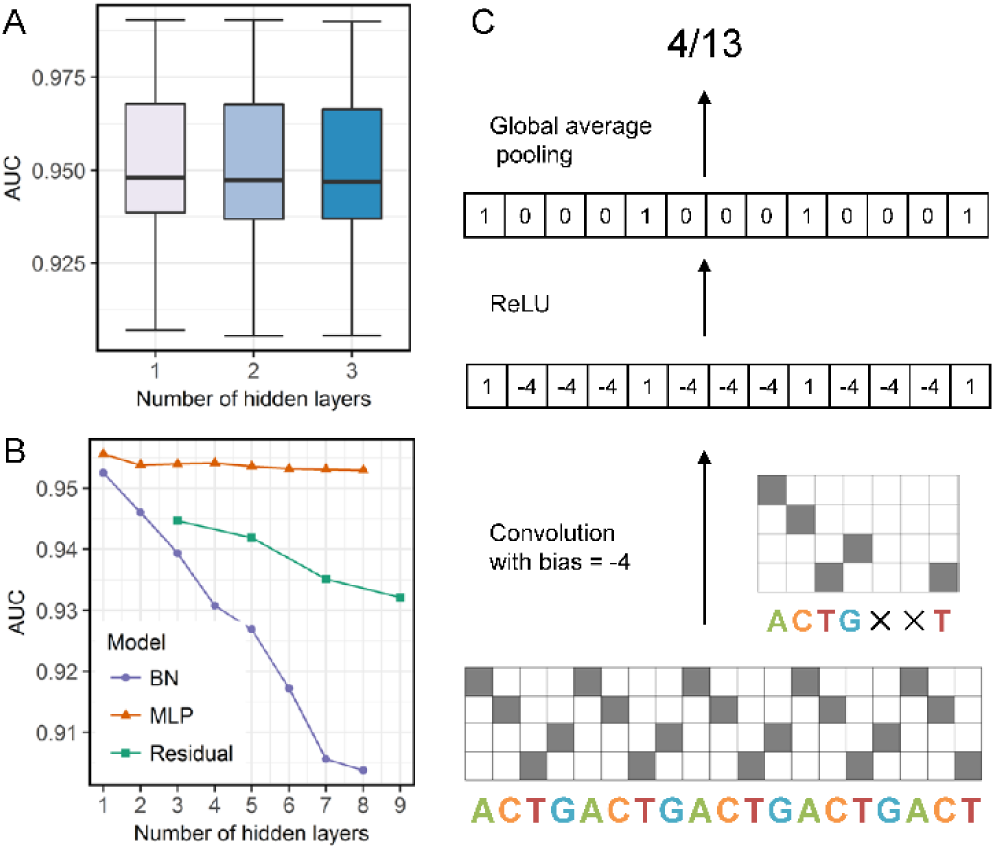
Exploration of advanced gkm-DNN and its connection with CNN. (A) Box plots of AUC for models with different hidden layers on 69 high-quality datasets. The bottom, top, and middle bands of the boxes indicate the 25th, 75th, and 50th percentiles, respectively. Whiskers extend to the most extreme data points no more than 1.5 interquartile range from the box. (B) AUCs of different models on CTCF dataset of K562 cell line. Multi-layer perception (MLP) represents our simple model. Batch normalization (BN) represents simple models with BN right before each ReLU activation. Residual nets (Residual) represents BN models with residual connections between successive hidden layers of odd number. (C) Counting the frequency of gapped k-mer is equivalent to performing a convolution operation with a global average pooling on the one-hot coding format of a DNA sequence.

Linking it with convolutional neural networks (CNN) may help to explain the phenomenon. First, a gapped k-mer can be transformed (e.g. ACTG××T) to a convolution kernel (Fig. 7C). Then, on the one-hot coding of DNA sequences, one can use the previous convolution kernel to convolve on the whole DNA sequence and then average the result (alternatively, the global average pooling layer). This is exactly equivalent to counting the frequency of that gapped k-mer.

In this framework, the gkm-fvs have several pros and cons. For advantages, the convolution step automatically extract high-level information of a DNA sequence without information loss and any time consuming learning. And the global average pooling operation greatly reduce the dimensions to make the computation feasible. However, the convolution kernels are not learned and a large num-ber of given convolution kernels (represented by gapped k-mers) are used as a compensation. The global average pooling layer destroy the location information, which may artificially create information bottleneck. As a result, we cannot guarantee whether the given gkm-fvs are enough to extract the desired information. In short, the gkm-fvs are high-level non-linear feature sets and con-tains few structured information. This explains why linear kernel SVM and relatively shallow neural networks works well using gkm-fvs as input.

## 4 Discussion

In this paper, we presented a flexible and scalable method gkm-DNN to achieve feature representation and applied it to a prediction task. We proposed a more concise version of gkm-fvs which significantly reduce the dimensions of gkm-fvs with few information loss. Then we implemented an efficient method to calculate the gkm-fv of a given sequence at the first time. We took the widely studied TFBS prediction problem as an illustrative example. We evaluated gkm-DNN and compared it with the state-of-the-art gkm-SVM using 467 small and 69 big datasets. We showed that gkm-DNN can not only overcome the potential limitations of the high dimensionality, colinearity and sparsity of gkm-fvs, but also make competitive performance compared to gkm-SVM using the same gkm-fvs.

One possible difficulty to use gkm-DNN is that one need to choose many hyper-parameters to train a model. The biggest factor affecting the performance is l and k for gkm-fvs. We recommend l = 7, k = 5 for typical cases. The hyper-parameters of training neural networks have diverse effects on different datasets (Supplementary Fig. S9A). However, if we only use the defaults rather than nine sets of hyper-parameters, the reductions of performance are very limited (Supplementary Figs. S9B and S9C). In conclusion, it is acceptable to use the well-chosen default hyper-parameters for gkm-DNN.

The main difficulty of using gkm-fvs is the calculation problem caused by its extremely high dimension. gkm-SVM, LS-GKM and gkm-DNN represent three directions of using gkm-fvs. gkm-SVM and LS-GKM use kernel tricks (SVM) while they optimize in different ways. gkm-SVM 2.0 adopts a new algorithm to calculate the kernel matrix in a faster way while LS-GKM combines LIBSVM framework to reduce the unnecessary calculation for the full kernel matrix [17], [26], [34]. So, gkm-SVM can use relative longer gapped k-mers by kernel tricks while it cannot handle large amount of samples. Although LS-GKM improves this [26], the running is still not efficient enough. On the other hand, gkm-DNN tries to directly use gkm-fvs as inputs without kernel tricks. It simplifies gkm-fvs from structure and can deal with large amount of data using relatively shorter gapped k-mers. For future work, how to more efficiently calculate the string kernel matrix or to further simplify the gkm-fvs may be two valuable directions.

More importantly, gkm-DNN directly uses gkm-fvs and should be viewed as a DNA feature extraction method. Since the concise gkm-fvs is a natural extension of lmer-fvs, many methods using lmer-fvs may benefit from using the concise gkm-fvs instead. On the other hand, with the rapid development of biotechnologies, large-scale sequence data and diverse types of other data are emerging. In this background, more and more methods try to combine different data types rather than using DNA sequence data alone [35], [36]. Therefore, integrating gkm-DNN with other data type to achieve more biologically specific tasks will be valuable in future.

In summary, it is easy to extend gkm-DNN due to its high flexibility and scalability. First, one can uses large amount of data with time scaling in proportion to the sample size. Second, gkm-DNN can also add other data sources as inputs using a computational graph, which is a directed acyclic graph representing the information flow (Supplementary Fig. S10). The hidden layers automatically extract the high-level features and allocate different weights for different data sources. Third, gkm-DNN can deal with a variety of outputs such as multi-label classification and regression (Supplementary Fig. S10). The only change is to use a proper activation function for the output layer and a proper loss function for training. Therefore, we can easily adapt gkm-DNN to many other prediction tasks such as transcription factor binding affinity prediction and enhancer recognition [35], [37].

## Acknowledgment

Shihua Zhang is the corresponding author of this paper. This work has been supported by the National Natural Science Foundation of China [No. 61422309, 61379092, 61621003 and 11661141019]; the Strategic Priority Research Program of the Chinese Academy of Sciences (CAS) [No. XDB13040600], the Key Research Program of the Chinese Academy of Sciences [No. KFZD-SW-219] and CAS Frontier Science Research Key Project for Top Young Scientist [No. QYZDB-SSW-SYS008].

